# Inhibition of *Campylobacter jejuni* biofilm formation by D-amino acids

**DOI:** 10.1101/2020.07.30.230045

**Authors:** Bassam A. Elgamoudi, Taha, Victoria Korolik

**Affiliations:** Institute for Glycomics, Griffith University, Gold Coast campus, QLD 4222, Australia

**Keywords:** D-amino acids, *Campylobacter jejuni*, Biofilm, Alanine racemase, CLSM, confocal laser scanning microscopy

## Abstract

The ability of bacterial pathogens to form biofilms is an important virulence mechanism in relation to its pathogenesis and transmission. Biofilms play a crucial role in survival in unfavourable environmental conditions, act as reservoirs of microbial contamination and antibiotic resistance. For intestinal pathogen *Campylobacter jejuni*, biofilms are considered to be a contributing factor in transmission through the food chain and currently, there are no known methods for intervention. Here we present an unconventional approach to reducing biofilm formation by *C. jejuni* by the application of D-amino acids (DAs), and L-amino acids (LAs). We found that DAs and not LAs, except L-alanine, reduced biofilm formation by up to 70%. The treatment of *C. jejuni* cells with DAs changed the biofilm architecture and reduced the appearance of amyloid-like fibrils. In addition, a mixture of DAs enhanced antimicrobial efficacy of D-Cycloserine (DCS) up to 32% as compared with DCS treatment alone. Unexpectedly, D-alanine was able to reverse the inhibitory effect of other DAs as well as that of DCS. Furthermore, L-alanine and D-tryptophan decreased transcript levels of peptidoglycan biosynthesis enzymes alanine racemase (*alr*) and D-alanine-D-alanine ligase (*ddlA*) while D-serine was only able to decrease the transcript levels of *alr*. Our findings suggest that a combination of DAs could reduce biofilm formation, viability and persistence of *C. jejuni* through dysregulation of *alr* and *ddlA.*

## Introduction

Human pathogen *Campylobacter jejuni* is a leading foodborne bacterial cause of diarrhoeal disease which, according to the World Health Organization (WHO), occurs annually in approximately 10% of the world’s population, including 200 million children (1, 2). Campylobacters are increasingly resistant to antibiotics which is enhanced by their ability to form biofilms (3–5). *C. jejuni,* in particular, is able to form mono- and mixed-culture biofilms *in vitro* and *in vivo* (6), which recognized as a contributing factor of *C*. *jejuni* transmission through the food chain where biofilms allow the cells to survive up to twice as long under atmospheric conditions and in water (7–9). Campylobacters exhibit intrinsic resistance to many antimicrobial agents such as cephalosporins, trimethoprim, sulfamethoxazole, rifampicin and vancomycin, and are listed in WHO list of priority pathogens for new antibiotics development (3, 4, 10–15). Biofilms are known to enhance antimicrobial resistance of many pathogens (3–5, 16); thus, unconventional approaches to controlling biofilms and improving the efficacy of currently used antibiotics are urgently needed. Recent investigations into potential antimicrobials include naturally occurring small molecules such as nitric oxide, fatty acids, and D-amino acids (DAs) (17–20). DAs showed an ability to disperse some bacterial biofilms *in vitro*, such as those formed by *Bacillus subtilis*, *Staphylococcus aureus*, *Enterococcus faecalis* and *Pseudomonas aeruginosa* (21–26). It is well documented that microorganisms preferentially utilize L-amino acids (LAs) over DAs (27, 28), yet naturally occurring DAs have been found in different environments, such as soil, as well as in human and animals tissues (27). In addition, many bacterial species secrete DAs in the stationary growth phase and when encased in biofilms. For example, *Vibrio cholerae* can produce D-methionine (D-met) and D-leucine (D-leu), while *B. subtilis* generates D-tyrosine (D-tyr) and D-phenylalanine (D-phe) which can accumulate at millimolar concentrations (29, 30). The ability of bacteria to produce DAs is proposed to be a mechanism for self-dispersal of aging biofilms, and DA production may also inhibit the growth of other bacteria during maturation of mixed biofilms. In a naturally occurring biofilms, DAs are found to be involved in the regulation of extracellular polymeric saccharide (EPS) production, for instance, D-tyr reduces the attachment of *B. subtilis, S. aureus* and *P. aeruginosa* to surfaces (22, 24, 31–33). Also, DAs can induce disassembly of matrix-associated amyloid fibrils that link the cells within the biofilm and contribute to the biofilm strength (34). Effective concentration of DAs required to inhibit the biofilm formation varies depending on bacterial strain and DAs concentration ranging between 3 μM and 10 mM (23, 33, 35, 36). It’s important to note that some DAs exhibit inhibitory or toxic effects on a number of bacterial species and can interfere with the activities of peptidases and proteases involved in cell wall synthesis, for example, D-met can be incorporated into the peptidoglycan (PG) of bacterial cell walls, causing morphological and structural damage (37).

DAs appear to be able to disrupt the biofilms via multiple mechanisms, offering an advantage to other biofilm dispersal agents which target a single process essential for biofilm formation, indicating that DAs could form basis for a potential antibiofilm agent.

Herein, we demonstrate that D-alanine (D-ala), L-alanine (L-ala), D-serine (D-ser), D-methionine (D-met), and D-tryptophan (D-trp) can inhibit and disperse biofilms formed by *C. jejuni* and *C. coli* and that it may be possible to use these DAs to enhance the efficacy of antibiotics such D-cycloserine. Also, we present evidence that DAS target alanine racemase (*alr*) in *C. jejuni,* which leads to the inhibition of growth and biofilm formation. This finding may be the key to understanding the mechanisms of DAs action and also could provide an alternative strategy to control *Campylobacter* spp transmission via the food chain.

## Results

### Effect of LAs and DAs on biofilm formation by *C. jejuni*

In order to investigate the effect of LAs and DAs on biofilm formation, different concentrations of LAs and DAs (0.1-100 mM) were tested for their ability to disrupt or disperse the *Campylobacter* biofilm. Two assays were applied, one to measure the percentage of biofilm inhibition (%) (Inhibition Assay) and the other to determine the effect on the dispersion of a formed biofilm (Dispersion Assay). Treatment of *C. jejuni* culture with DAs showed significant inhibitory effect (*P* < 0.001) on biofilm formation. Prescreening of individual LAs and DAs identified four (D-ala, D-met, D-ser, and D-trp) that had a potent ability to inhibit biofilm formation by *C. jejuni* (Fig 1). In contrast, the L-form of those amino acids, except L-ala, had no inhibitory effect, and some of them, L-met and L-trp, significantly increased biofilm formation.

**Fig 1.**
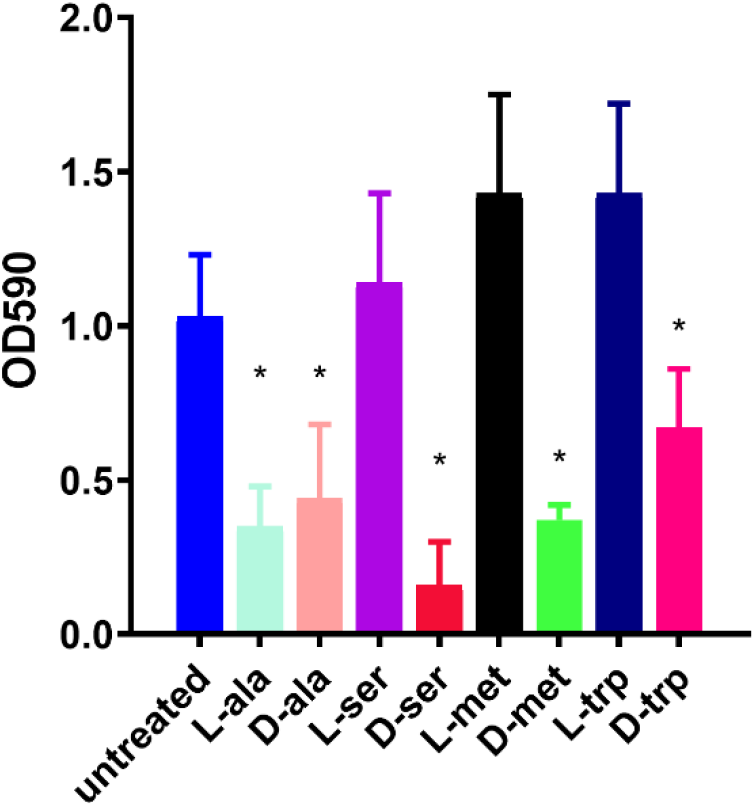
Effect of 100 mM DAs and LAs on *C. jejuni* 11168-O biofilm. Inhibition of biofilm formation in the presence of 100 mM of; L-alanine (L-ala), D-alanine (D-ala), L-serine (L-ser), D-serine (D-ser), L-methionine (L-met), D-methionine (D-met), L-tryptophan (L-trp), or D-tryptophan (D-trp). The asterisk (*) indicates a statistically significant difference using the unpaired Student’s t-test, p<0.05.

The DAs had a strong inhibitory effect on biofilm formation by *C. jejuni* at 10 mM concentration, with 48% inhibition for D-trp, while D-ala reduced biofilm formation by 28%. Interestingly, 50 mM L-ala reduced biofilm by up to 63% as compared to 45% by D-ala at same concertation (Fig 2). DAs had a disruptive effect on the existing biofilm where D-ser had the most significant effect (*P* < 0.001) on formed biofilm disruption, up to 71%, at 50 mM (Fig 2), and the addition of 10 mM D-trp led to 42% disruption of formed biofilm. Based on the results of DAs inhibitory and dispersal activities, the concentration between 5 to 10 mM was chosen for all subsequent assays.

**Fig 2.**
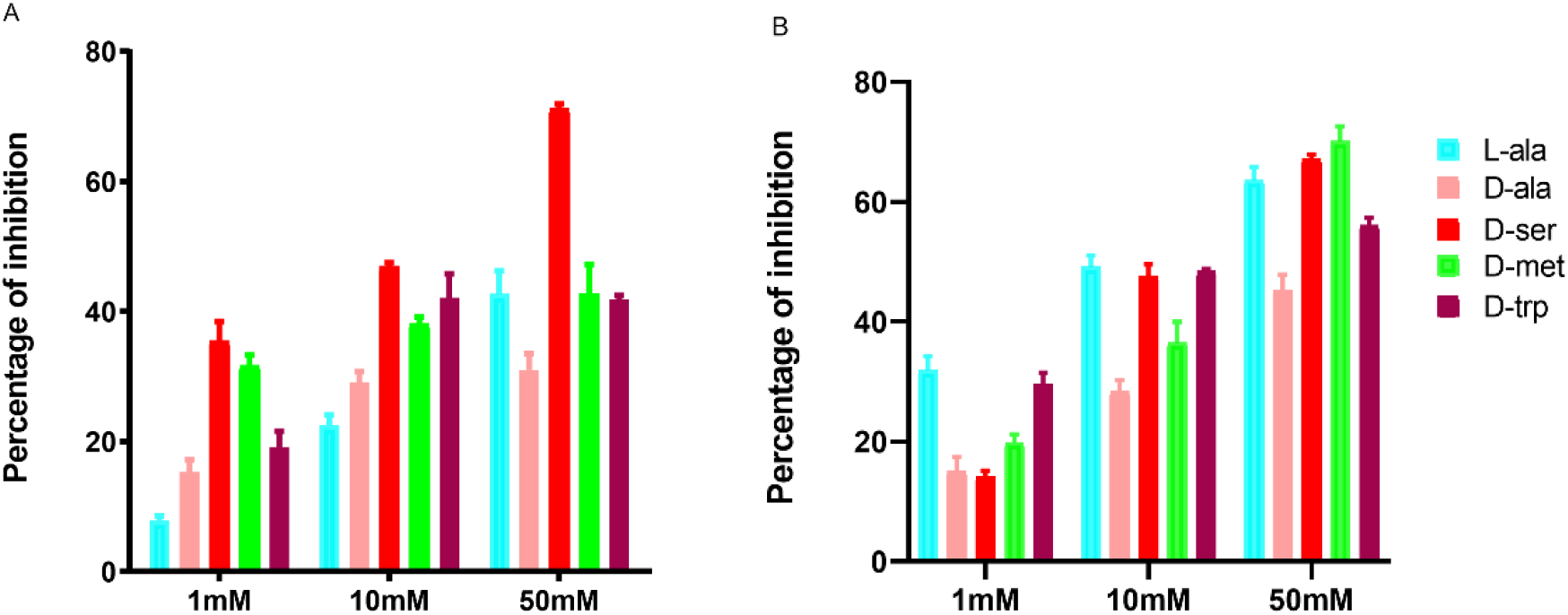
Inhibition and dispersion response of *C. jejuni* 11168-O biofilms in the presence of LAs and DAs at different concentrations. A) Dispersion of the existing biofilm induced by different concentrations of LAs and DAs. B) Inhibition of biofilm formation by different concentrations of LAs and DAs.

In order to elucidate strain-specific responses, *C. jejuni* 11168-O*, C. jejuni* 81-176, and *C. coli* NCTC 11366, were used to confirm the inhibitory effect of D-ala, D-ser, D-met, and D-trp at 10 mM. The effect of DAs on biofilm formation was strain-dependent, where D-ser and D-trp had the greatest inhibitory effect on biofilm formation by 11168-O, D-ala and D-met were most effective against 81-176, and *C. coli* (Fig 3).

**Fig 3.**
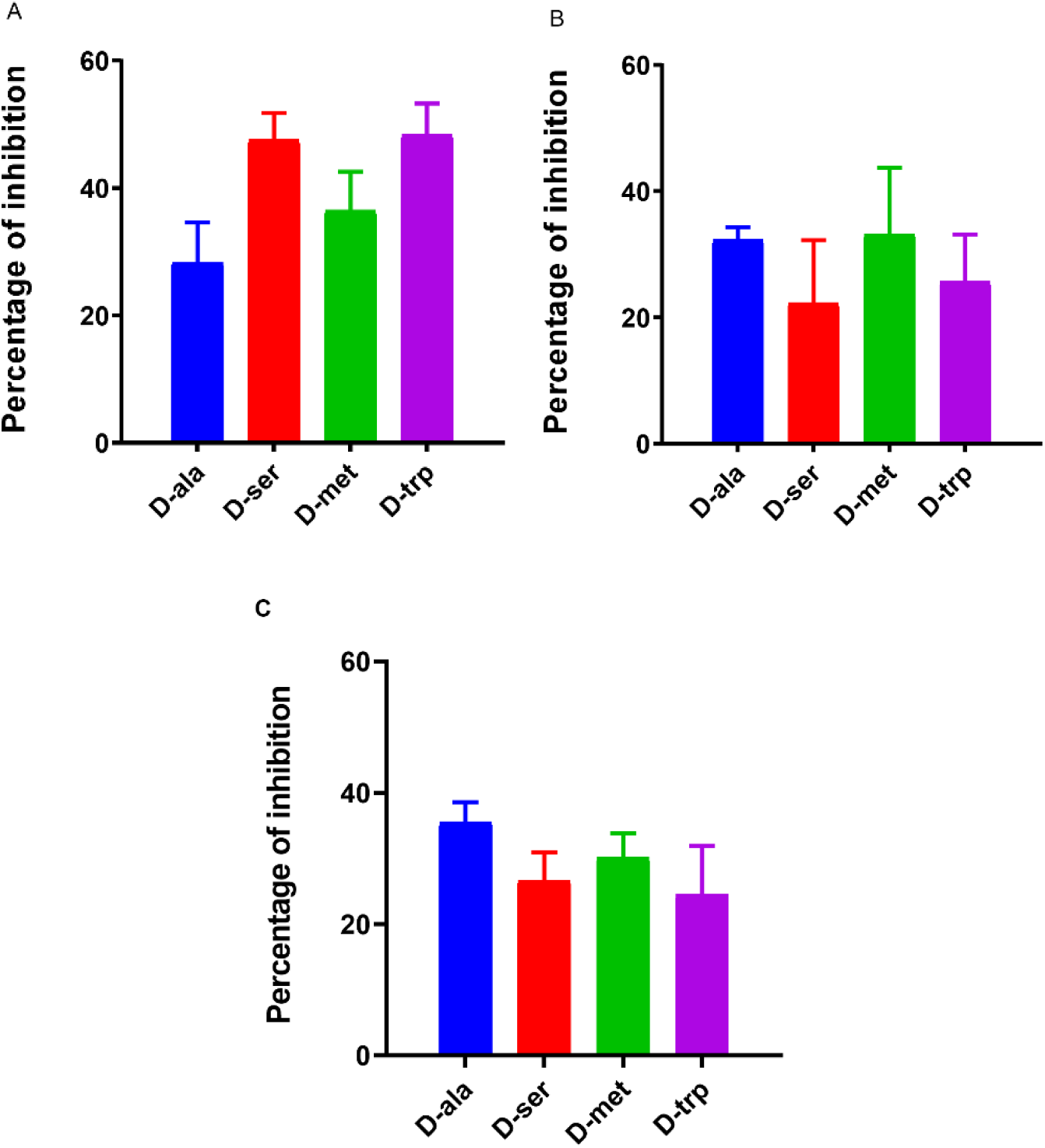
Quantitative analysis of biofilm inhibition of A) *C. jejuni* 11168-O, B*) C. jejuni* 81-176, and C) *C. coli* NCTC 11366 in the presence of 10 mM of DAs.

The equimolar mixture of DAs and LAs (1:1) showed ≥ 40% inhibition of *C. jejuni* 11168-O biofilm formation (Fig 4). The mixture of the four amino acids, L-ala, D-met, D-ser, D-trp (5:5:2:5 mM), was more potent, with up to 49% inhibition of biofilm formation; however, the addition of D-ala to D-ser decreased the inhibitory effect (Fig 4).

**Fig 4.**
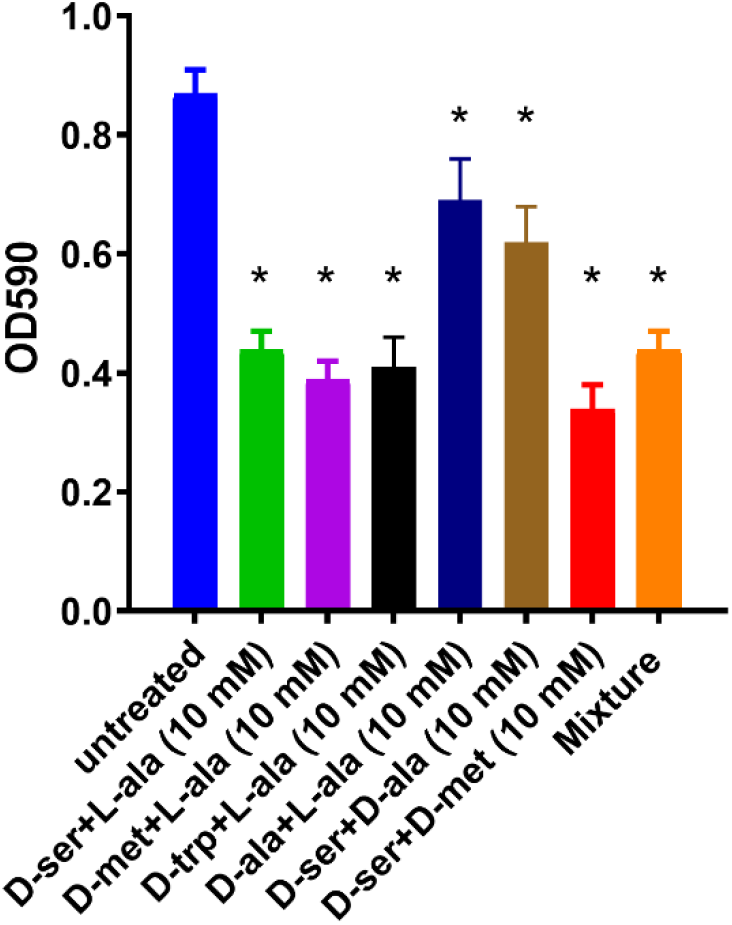
Effect of the equimolar mixture of DAs and LAs on *C. jejuni* 11168-O biofilm. The asterisk (*) indicates a statistically significant difference using the unpaired Student’s t-test, p<0.05.

### Microscopic characterization of the dispersion effect of DAs on biofilm formation

Microscopic examination of formed biofilms, treated with individual DAS, by confocal microscopy demonstrated a significant reduction in biofilm formation compared to that of untreated controls (Fig 5). The disassembly of the amyloid fibrils, which connect the cells within the structure of *C. jejuni* biofilms, can also be observed (Fig 6).

**Fig 5.**
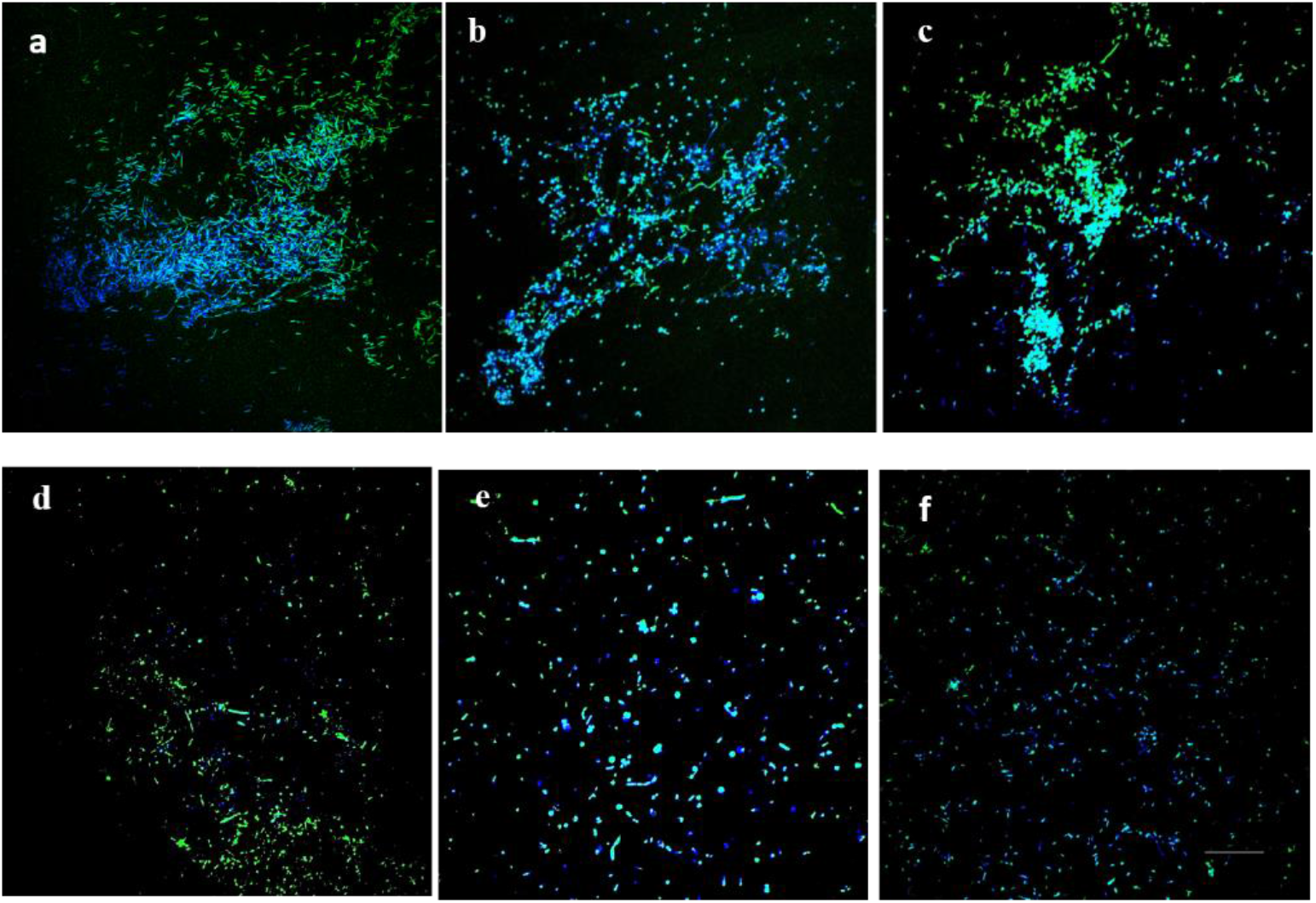
Confocal scanning laser microscopy images of *C. jejuni* 11168-O biofilm in presence of 25 mM of DAs. *C. jejuni* biofilm at 48h, imaged using dual fluorescence labelling by CLSM. a) Untreated, b) D-ala, c) L-ala, d) D-ser, e) D-met, f) D-trp. Cells were stained with 4′,6-diamidino-2-phenylindole (DAPI, blue) and amyloid fibrils by Thioflavin T (ThioT, green) (Scale bar= 20μm).

**Fig 6.**
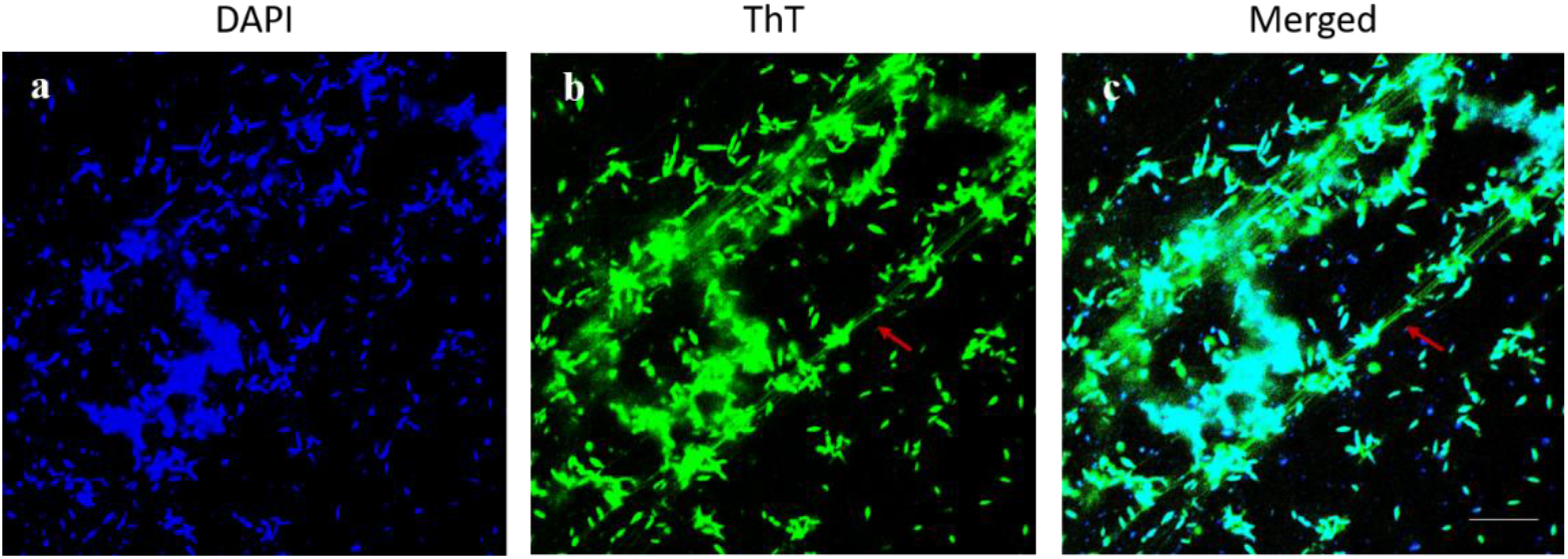
The mature biofilm of *C. jejuni* 11168-O and amyloid-like fibres. *C. jejuni* biofilm imaged using dual fluorescence labelling by CLSM. Red arrow indicates for amyloid-like *fibrils* (ThioT, green) and bacterial cells within the biofilm (DAPI, blue). (Scale bar= 10μm).

### Expression level of *alr* and *ddlA* in the presence of LAs and DAs

In order to interrogate the mechanism of inhibitory action of DAs and L-ala, the expression of *C. jejuni* PG biosynthesis enzymes alanine racemase (*alr*) and D-Ala-D-Ala ligase (*ddlA*) in the presence and absence of DAs and LAs were examined. The relative expression of *ddl* and *alr* was downregulated by 1.25 to 4-fold below the cut-off level, respectively, following treatment of cells with 25 mM of L-ala (Table 1). In contrast, 25 mM of D-ala upregulated the expression of *ddl* by 10-fold and *alr* by 38-fold. Treatment of cells with 25 mM D-trp downregulated the expression level of *ddl* by 1.65-fold and *alr* by 3-fold whereas D-ser (25 mM) downregulated the expression of *alr* by 2.92-fold and upregulated *ddl* by 2.58-fold. No significant effect on the expression of *alr* and *ddl* was observed following treatment with D-met (Table 1). Interestingly, treatment of cells with D-Cycloserine (DCS) (10ng/ml), as a positive control, had a greater effect, downregulating the expression of *ddlA* with a 7-fold change as compared to 2.85-fold change for *alr*. No loss of cell viability could be detected after 2-h exposure to DAs or DCS.

**Table 1.** Analysis of the relative expression of *alr* and *ddlA* genes in the present of LAs and DAs by real-time PCR (qRT-PCR). The relative expression of *alr* and *ddl* genes after incubation of *C. jejuni* 11168-O cells with 25 mM of LAs and DAs for 2 hrs.

| Gene name | Fold change |  |  |  |  |  |  |
| --- | --- | --- | --- | --- | --- | --- | --- |
|  | upregulated |  |  | downregulated |  |  |  |
|  | D-ala | D-ser | D-met | L-ala | D-ser | D-trp | DCS |
| *alr* | 38±7 | - | - | 4.18±0.3 | 2.92±0.2 | 1.64±0.3 | 2.85±0.2 |
| *ddlA* | 10±2 | 2.58±0.6 | - | 1.25±0.1 | - | 3.42±0.4 | 7.15±0.2 |

### D-Ala can reverse the inhibitory effect of DAs and DCS

D-ala has been reported to reverse the antimicrobial efficacy of DCS in *Mycobacterium* spp (38, 39). Considering that the MIC range of DCS for *Campylobacter* spp reported to be between 0.25 μg/ml-4 μg/ml (40), we tested the effect of sub-inhibitory concentration of 50ng/ml DCS on *C. jejuni* cells and determined that DCS can reduce *C. jejuni* growth and biofilm formation by up to 76% (Fig 7). Furthermore, this effect can be reversed by increasing the concentration of D-ala from 10 mM to 50mM (Fig 7). Combining D-ala with other DAs also decreased the inhibition of biofilm formation. In contrast, a combination of DAs with DCS increased the efficacy of DCS at 10 ng/ml by 32% as compared with DCS treatment alone (Fig 8).

**Fig 7.**
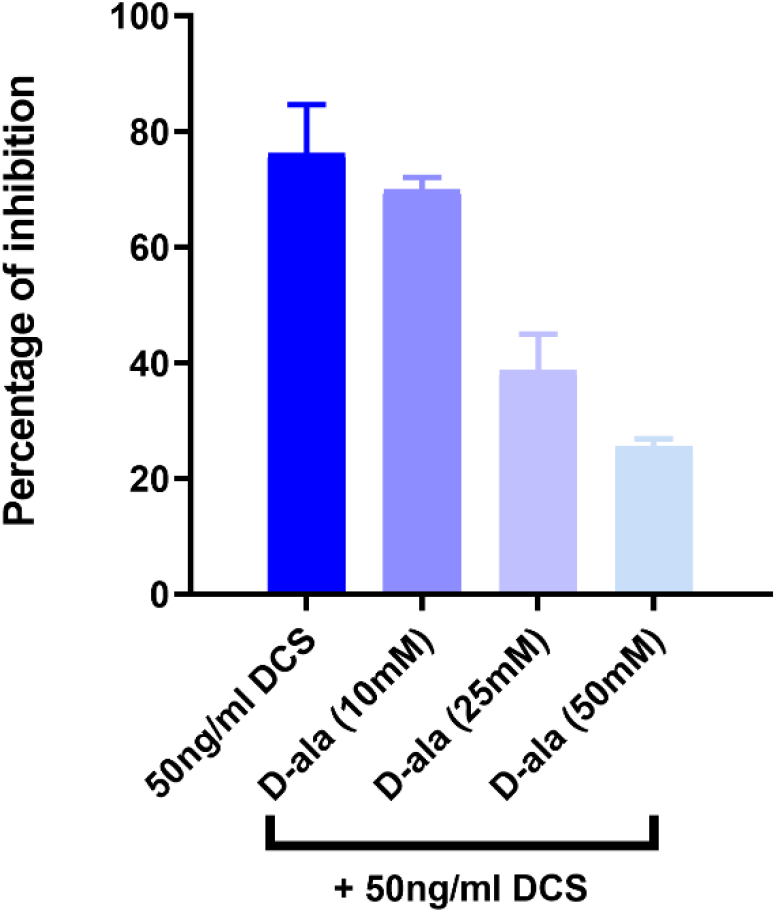
Reversal of DCS growth inhibition by D-alanine at different concentrations in *C. jejuni* 11168-O.

**Fig 8.**
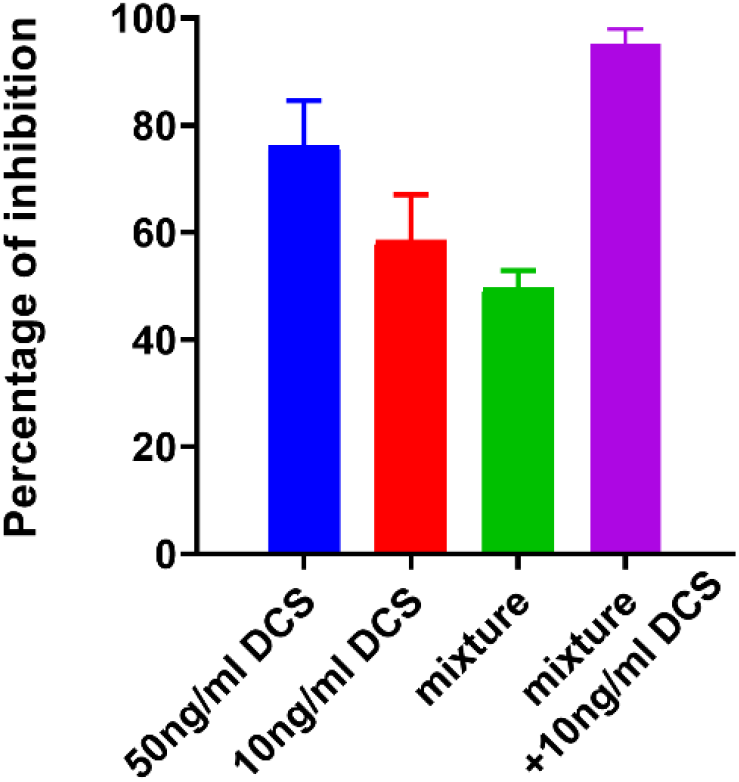
Effect of DCS on *C. jejuni* 11168-O biofilm when combined with L-ala, D-ser, D-met, D-trp (5:5:2:5 mM).

## Discussion

This study describes the identification of specific small, naturally occurring molecules, DAs, which are highly effective in preventing and disrupting *C. jejuni* biofilms, in concert with that previously shown for *B. subtilis, S. aureus* and *P. aeruginosa* (24, 36, 41). While D-met and D-trp are able to inhibit the biofilm formation of *C. jejuni*, L-form of those amino acids significantly increased biofilm formation. It is possible that *C. jejuni* is able to catabolize L-form of those amino acids (42), which promotes bacterial growth, and consequently formation of the biofilm. This is consistent with the previous report of *B. subtilis* growth inhibition by D-form of Tyr, Leu, and Trp, and the L-form of those amino acids counteracting this effect (23). The effect of DAs on inhibition and dispersal of *C. jejuni* biofilms showed a concentration-dependent response, with D-ser, D-met and D-trp being most effective in inhibition and dispersion of the biofilm. We observed that D-met, and D-trp, have a significant dispersive effect on biofilms at concentrations of ≥10 mM, similar to that observed for *S. aureus* and *P. aeruginosa* (43). It’s important to note that, the inhibitory effect on the growth of *C. jejuni* by DAs, except D-met, could be reversed by D-ala, similar to that observed for *B. subtilis*, *M. tuberculosis* and *Escherichia coli* (38, 39, 44, 45).

Microscopic analysis confirmed the effect of DAs on biofilm formation of *C. jejuni,* and particularly, the formation of amyloid-like fibrils within the biofilm matrix. Matrix-associated amyloid fibrils had been previously reported to form a part of C. jejuni biofilm (46), and similar DA-induced disassembly of matrix-associated amyloid fibers of *B. subtilis* biofilm, had been proposed as a biofilm-dispersal mechanism (34, 41). Together, these data allow us to speculate that the ability of DAs to promote the dispersal of formed C. *jejuni* biofilms, could involve the triggering the disassembly of matrix-associated amyloid fibrils. While the mechanisms of antimicrobial and antibiofilm action of DAs, particularly, D-ser, D-met, and D-trp, are not fully understood, DAs effect on *C. jejuni* growth and biofilm formation may be similar to that for *Alcaligenes faecalis,* where D-met incorporates into PG, causing morphological and structural damage to the cell wall (30, 37, 47), and consequently suppresses bacterial growth. To explore that possibility, we interrogated the effect of DAs and LAs on the expression level of two genes in *C. jejuni*; alanine racemase (*alr*) (*Cj0905c*), and D-Ala-D-Ala ligase (*ddlA*) (*Cj0798c*) (48, 49). Both genes are encoding enzymes involved in an important step in D-Ala metabolism (44, 50), which is essential for the synthesis of PG of the bacterial cell wall (45, 51, 52). Two main reactions are involved in this process, first the conversion of L-Ala to D-Ala by alanine racemase (alr), and the formation of D-alanyl–D-alanine dipeptide (D-Ala-D-Ala) from D-Ala by D-alanine–D-alanine ligase (ddl) (53). RT-PCR data shows that DCS able to reduce both *C. jejuni alr* and *ddlA* expression levels, similarly to L-ala, and D-trp. Interestingly, D-ser reduced *alr* expression levels, but not that of *ddlA*, suggesting that *ddlA* may not be the primary target for D-ser or DCS in *C. jejuni.* Furthermore, the ability of D-ala to reverse the inhibitory effect of DCS and D-ser suggests that the inhibitory effect of DCS and D-ser on *C. jejuni* can be mediated through inhibition of *alr* alone. In contrast, in *M. tuberculosis,* both *alr* and *ddl* were reported to be the primary targets of DCS (39), and *S. Halouska, et al. (54)* suggested that *ddl* ay be a primary target of DCS, rather than *alr*.

It is interesting to note that bacterial PG dipeptide D-Ala-D-Ala, which is generated by D-Ala-D-Ala ligase (*ddlA*), is the usual target for vancomycin, but in *C. jejuni,* PG contains D-Alanyl-D-Lactate (D-Ala-D-Lac) termini resulting in reduced efficacy of vancomycin by up to 1,000-fold. Substitution by D-alanyl-D-serine (D-Ala-D-ser) termini reduces the efficacy of this antibiotic by up to 7-fold (4, 55–58). This further suggests that *alr* and not *ddlA,* is likely to be the primary target for D-ser and DCS in *C. jejuni,*.

Our results suggest that DAs might have a promising application in enhancing the activity antibiotics where the combination of DAs with DCS, synergistically increased the ability of DCS to inhibit *C. jejuni* biofilm formation and growth. The enhancement of DCS efficacy with DAs is likely to lower minimal dose requirement, which would consequently reduce the drug toxicity. DAs had also been reported to enhance the effectiveness for colistin and ciprofloxacin, when used against biofilms of *P. aeruginosa*, and rifampin used against biofilms of clinical isolates of *S. aureus* (43).

To summarize, this study suggests that (i) DAs show the inhibitory effect at millimolar concentrations on biofilm formation by C. *jejuni*; (ii) DAs can trigger C. *jejuni* biofilm-disassembly; (iii) a combination of DAS can enhance the efficacy of DSC, (iv) DAs inhibit growth and biofilm formation of *C. jejuni* by repressing the expression of *alr.* The data described here contribute to the understanding of the mechanisms involved in biofilm dispersion and inform on identification of potential antimicrobial drug targets.

## Materials and Methods

### *C. jejuni* strains and growth conditions

Bacterial strains used in this study were *C. jejuni* 11168-O (courtesy of Prof. D. G. Newell, United Kingdom), *C. jejuni* 81-176 (courtesy of Prof. Christine Szymanski, University of Alberta, Alberta), and *C. coli* NCTC 11366 (Griffith University culture collection, Australia). Cells were grown at 42°C microaerobically (85% N_2_, 10% CO_2_ and 5% O_2_) on Mueller-Hinton agar (MHA) and in Mueller-Hinton broth (MHB), supplemented with Trimethoprim (5 μg ml^−1^) and Vancomycin (10 μg ml^−1^) (TV) (Sigma).

### Chemical and reagents used in this study

L-alanine (L-ala), D-alanine (D-ala), L-serine (L-ser), D-serine (D-ser), L-methionine (L-met), D-methionine (D-met), L-tryptophan (L-trp), D-tryptophan (D-trp) D- cycloserine were from Sigma-Aldrich. Individual stock solutions of 100 mM of DAs in Phosphate-buffered saline (PBS) (PH 7.2).

### Biofilm formation and dispersion assays

Overnight cultures of *C. jejuni* strains were diluted to an OD_600_ of 0.05, and 2 mL of cell suspension was placed into 24-wells flat-bottom polystyrene tissue culture plates (Geiner Bio-One). Different concentrations of DAs (1-100mM) were added directly to the culture in the wells and incubated at 42°C under microaeroaerobic conditions for 48 hours. For dispersion assay, *C. jejuni* cells were grown as described above, except no DAs were added. Then PBS containing the appropriate concentration of DAs (0.1-100 mM) was added to the wells and plates incubated for further 24 hrs. For crystal violet staining, plates were rinsed with water once (gently) and dried at 55°C for 30 minutes and stained using modified crystal violet staining method as described previously (59). Data are representative of three independent experiments, and values are expressed in presented as Mean± S.D.

### RNA extraction, cDNA synthesis and RT-qPCR of Alanine racemase (*alr*), D-alanine-D-alanine ligase (*ddlA*)

*C. jejuni* 11168-O cells were grown overnight microaerobically in MHB at 42°C. Cells were collected by centrifuging at 4000 rpm for 15 minutes. The pellets were suspended in MHB and OD_600_ adjusted to 1 (~3×10^9^ cells/ml) and subsequently challenged with (1) 25 mM of L-ala, (2)25 mM of D-ala, (3) 25 mM of D-ser, (4) 25 mM of D-met or, (5) 25 mM of D-trp for 2-h; (5) 10ng/ml of DCS (below MIC which 250 ng/ml) was used as control. The bacterial survival was confirmed by viable cells counts after 2-h. Then, cells were collected by centrifugation at 4000 rpm for 15 minutes and pellets used for RNA extraction by RNeasy kit according to the manufacturer’s protocol (Bioline). cDNA synthesis and RT-qPCR was performed as previously described (60). The following primers sets were used: *alr* (Cj0905c) forward 3-AGCCAAAAATTTAGGAGTTT-5 and *alr* reverse 5-GAGGACGATGTGATAGTATT-3, *ddl* (Cj0798c) forward 3-TTATTTTTTGTGATGAAGAAAGAA-5 and *sdl* reverse 5-GAGTTCTTTTTCTTTTTTATAAGC-3. A *gryA* gene was used as a housekeeping control gene, using the primers, *gryA* forward 3-CCACTGGTGGTGAAGAAAATTTA-5 and *gryA* reverse 5-AGCATTTTACCTTGTGTGCTTAC-3. Relative *n*-fold changes in the transcription of the examined genes between the treated and non-treated samples were calculated using the relative quantification (RQ), also known as 2^−ΔΔCT^ method, where ΔΔ*C_T_*= Δ*C_T_* (treated sample) − Δ*C_T_* (untreated sample), Δ*C_T_* = *C_T_* (target gene) − *C_T_* (*gyrA*), and *C_T_* is the threshold cycle value for the amplified gene. The fold change due to treatment was calculated as −1/2^−ΔΔCT^ (61, 62). The data are presented as Mean± S.D and were calculated from triplicate cultures and are representative of three independent experiments.

### Confocal laser scanning microscopy

Overnight cultures of *C. jejuni* cells were diluted to an OD_600_ of 0.05, and 3 mL of each sample was placed into duplicate wells of a 6-well flat-bottom polystyrene tissue culture plate containing a glass coverslip to enable the formation of biofilm (Geiner Bio-One). 25 mM of LAs and DAs were added directly to the wells, and then the plates were incubated at 42°C microaerobically for 48 hours. After the incubation, MH broth was removed, and the wells were gently washed with PBS solution twice to remove planktonic cells. The coverslips were carefully removed by using sterile needle and forceps to new 6-well plates and fixed using 5% formaldehyde solution for 1 h at room temperature. Then, the coverslips were gently washed with 2 mL of PBS and prepared for staining with fluorescent dyes.

### Staining of *C. jejuni* cells

The fluorescent DNA-binding stain DAPI (Sigma Aldrich) was used to visualise cell distribution as described previously (63). Thioflavin T (ThT) at 20 μM was then used to treat the coverslips for 30 minutes. ThT emits green fluorescence upon binding to cellulose or amyloids (64, 65). The coverslips then were mounted on glass slides using the mounting medium (Ibidi GmbH, Martinsried, Germany) and sealed with transparent nail varnish. Microscopy (Nikon A1R+) (Griffith University) was performed with two coverslips per sample from at least two separate experiments. All images were processed using ImageJ analysis software version 1.5i (National Institutes of Health, Besthda, Maryland).

### Statistical analysis

The statistical analyses performed using GraphPad Prism, version 6.00 (for Windows; GraphPad Software) to calculate statistically significant differences when *P*- value by applied Student’s *t*-test.

## Author Contributions

VK and BE conceived and designed the study; BE and Taha performed the experiments; VK, and BE, analyzed the data and prepared the manuscript. All authors reviewed the manuscript.

## Acknowledgements

The authors thank Neeraj Bhatnagar, Michael Hadjistyllis for technical assistance.

